# Homeostatic plasticity and external input shape neural network dynamics

**DOI:** 10.1101/362152

**Authors:** Johannes Zierenberg, Jens Wilting, Viola Priesemann

**Affiliations:** Max Planck Institute for Dynamics and Self-Organization, Am Fassberg 17, 37077 Göttingen, Germany; Bernstein Center for Computational Neuroscience, Am Fassberg 17, 37077 Göttingen, Germany

## Abstract

*In vitro* and *in vivo* spiking activity clearly differ. Whereas networks *in vitro* develop strong bursts separated by periods of very little spiking activity, *in vivo* cortical networks show continuous activity. This is puzzling considering that both networks presumably share similar single-neuron dynamics and plasticity rules. We propose that the defining difference between *in vitro* and *in vivo* dynamics is the strength of external input. *In vitro*, networks are virtually isolated, whereas *in vivo* every brain area receives continuous input. We analyze a model of spiking neurons in which the input strength, mediated by spike rate homeostasis, determines the characteristics of the dynamical state. In more detail, our analytical and numerical results on various network topologies show consistently that under increasing input, homeostatic plasticity generates distinct dynamic states, from bursting, to close-to-critical, reverberating and irregular states. This implies that the dynamic state of a neural network is not fixed but can readily adapt to the input strengths. Indeed, our results match experimental spike recordings *in vitro* and *in vivo*: the *in vitro* bursting behavior is consistent with a state generated by very low network input (< 0.1%), whereas *in vivo* activity suggests that on the order of 1% recorded spikes are input-driven, resulting in reverberating dynamics. Importantly, this predicts that one can abolish the ubiquitous bursts of *in vitro* preparations, and instead impose dynamics comparable to *in vivo* activity by exposing the system to weak long-term stimulation, thereby opening new paths to establish an *in vivo*-like assay *in vitro* for basic as well as neurological studies.

## I. INTRODUCTION

Collective spiking activity clearly differs between *in vitro* cultures and *in vivo* cortical networks (see examples in Fig. 1). Cultures *in vitro* typically exhibit stretches of very little spiking activity, interrupted by strong bursts of highly synchronized or coherent activity [1–7]. In contrast, spiking activity recorded from cortex in awake animals *in vivo* lacks such pauses, and instead shows continuous, fluctuating activity. These fluctuations show a dominant autocorrelation time that was proposed to increase hierarchically across cerebral cortex, from sensory to frontal areas [8]. Moreover, depending on experimental details such as brain area, species and vigilance state, one also observes evidence for asynchronous-irregular (AI) dynamics [9, 10], oscillations [11–13], or strong fluctuations associated with criticality, bistability or up-and-down states [14–20]. These states differ not only in strength and structure of fluctuations, but also in synchrony among neurons, from uncorrelated to fully synchronized spiking. The observation of such a vast range of dynamic states is puzzling, considering that the dynamics of all networks presumably originate from similar single-neuron physiology and plasticity mechanisms.

**FIG. 1.**
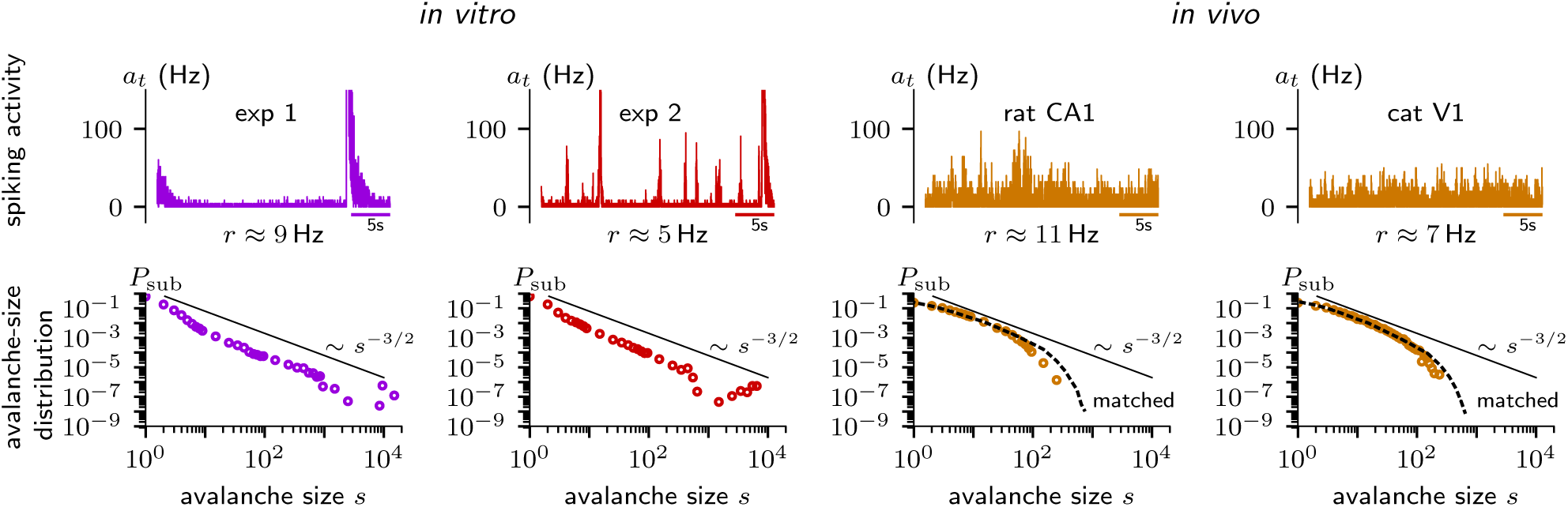
Examples of dynamic states observed in experiments. *In vitro* spike recordings are from cultures of dissociated rat cortical neurons [4]. *In vivo* recordings are from right dorsal hippocampus in an awake rat during an open field task [73] and from primary visual cortex in a mildly anesthetized cat [76]. Top row shows population spiking activity (Appendix B-1) from 30-60 single or multi-units (Δ*t* = 4 ms) with average neural firing rate *r*; bottom row shows subsampled avalanche-size distributions (Appendix B-2). Solid lines indicate the power-law behavior *s*^−3/2^ expected for a critical branching process. Dashed lines correspond to distributions obtained from branching networks matched to the experiments of rat CA1 (*τ* = 2 s, *r*^∗^ = 11 Hz, *h* = 5.5 × 10^−3^ Hz, *n* = 31, *N* = 104, Δ*t* = 1 ms, *τ*_hp_ = 10^5^ s) and cat V1 (*τ* = 0.2 s, *r*^∗^ = 7 Hz, *h* = 3.5 × 10^−2^ Hz, *n* = 50, *N* = 104, Δ*t* = 1 ms, *τ*_hp_ = 10^5^ s). For details and a definition of parameters see Appendix B-4.

One particular plasticity mechanism that regulates neural activity on a long time scale is homeostatic plasticity [21–26]. Homeostatic plasticity can be implemented by a number of physiological candidate mechanisms, such as redistribution of synaptic efficacy [27, 28], synaptic scaling [21–23, 29], adaptation of membrane excitability [25, 30], or through interactions with glial cells [31, 32]. All these mechanisms have in common that they implement a slow negative feedback-loop in order to maintain a certain target spike rate and stabilize network dynamics. In general, they reduce (increase) excitatory synaptic strength or neural excitability if the spike rate is above (below) a target rate, allowing compensation against a potentially unconstrained positive feedback-loop through Hebbian-type plasticity [33–39]. Recent results highlight the involvement of homeostatic plasticity in generating robust yet complex activity dynamics in recurrent networks [40–42].

To understand the physiological mechanisms behind this large set of dynamic states, different model networks have been proposed that reproduce one or a set of states. To name a few examples, deafferentiation in combination with homeostatic scaling can generate bursts [43]; the interplay between excitation and inhibition may lead to oscillations, synchronous-regular activity, or asynchronous-irregular activity [44–47], where switching between dynamic states can be induced by varying the input [47–49]; synaptic facilitation and depression promote regular and irregular network dynamics [50–52]; plasticity at inhibitory synapses can stabilize irregular dynamics [53, 54], whereas specific types of structural [55–57] or synaptic [51, 52, 58–66] plasticity foster strong temporal fluctuations characteristic for a critical state; last but not least, homeostasis is necessary to achieve stable dynamics in recurrent networks with spike-timing dependent plasticity (STDP) or Hebbian-type synaptic plasticity (e.g. Refs. [38, 67–71]). Overall, the dynamic state depends on all aspects: single-neuron properties, synaptic mechanisms, network topology, plasticity rules, and input characteristics. Recalling that the single-neuron properties, synaptic mechanisms, as well as plasticity rules are pre-sumably very similar *in vitro* and *in vivo*, these factors are unlikely to explain the observed differences.

In this study, we propose that the input strength is the defining difference between *in vitro* and *in vivo* dynamics. *In vitro* systems are completely isolated, whereas *in vivo* networks receive continuous input from sensory modalities and other brain areas. Under these different conditions, we propose that homeostatic plasticity is a sufficient mechanism to promote self-organization to a diverse set of dynamic states by mediating the interplay between external input rate and neural target spike rate. Treating the external input as a control parameter in our theoretical framework, allows us to alter the network dynamics from bursting, to fluctuating, to irregular. Thereby, our framework offers testable predictions for the emergence of characteristic but distinct network activity *in vitro* and *in vivo*.

Based on our theory, we derive explicit experimental predictions and implications: (1) The direct relation between dynamic state, spike rate and input rate enables us to quantify the amount of input the neural network receives, e.g., in mildly anesthetized cat V1, we estimate an input rate of 𝓞(0.01 Hz/neuron). (2) This implies that about 2% of cortical activity in cat V1 are imposed by the input, whereas 98% are generated by recurrent activation from within the network. (3) Our results suggest that one can alter the dynamic state of an experimental preparation by altering the input strength. Importantly, we predict for *in vitro* cultures that increasing the input rate to about 𝓞(0.01 Hz/neuron) would be sufficient to induce *in vivo*-like dynamics.

## II. EXPERIMENTAL OBSERVATIONS

To demonstrate characteristic neural activity *in vitro* and *in vivo*, we analyzed exemplary recordings of spiking activity. Data sets included cultures of dissociated cortical neurons [4, 72], as well as hippocampus of foraging rats [73, 74] and visual cortex of mildly anesthetized cats [75, 76] (see Appendix A for details). Note that all preparations were inevitably subsampled, as spikes were recorded only from a small number of all neurons. For illustrative purposes, we focus on the average (subsampled) spiking activity *a_t_* and the (subsampled) avalanche-size distribution *P*_sub_ (see Appendix B for details).

The spiking activity *in vitro* shows bursting behavior (Fig. 1), i.e., stretches of very low activity interrupted by periods of synchronized activity. The subsampled avalanche-size distributions *P*_sub_(*s*) exhibits partial power-law behavior resembling *P* (*s*) ~ *s*^−3/2^ as expected from a critical branching process [77], and conjectured for the synchronous-irregular regime [78]. However, in addition *P*_sub_(*s*) also shows a peak at large avalanche sizes, which may indicate either finite-size effects, supercriticality, or characteristic bursts [79].

In contrast, the spiking activity *in vivo* shows fluctuating dynamics (Fig. 1). These have been described as reverberating dynamics, a dynamic state between critical and irregular dynamics [80], characterized by a finite autocorrelation time of a few hundred milliseconds. The subsampled avalanche-size distributions *P*_sub_(*s*) can be approximated by a power-law for small *s* but show a clear exponential tail. The tails indicate slightly subcritical dynamics [81], especially because deviations in the tails are amplified under subsampling [15, 16, 79].

In sum, the spiking activity and the corresponding avalanche-size distributions clearly differ between *in vitro* and *in vivo* recordings. Remarkably, however, the average neural firing rate *r* is similar across the different experimental setups.

## III. MODEL

To investigate the differences between *in vitro* and *in vivo*, we make use of a branching network, which approximates properties of neural activity propagation. We extend the branching network by a negative feedback, which approximates homeostatic plasticity.

### A. Branching network

In the brain, neurons communicate by sending spikes. The receiving neuron integrates its input, and if the membrane potential crosses a certain threshold, this neuron fires a spike itself. As long as a neuron does not fire, its time-varying membrane potential can be considered to fluctuate around some resting potential. In the following, we approximate the complex time-resolved process of action potential generation and transmission in a stochastic neural model with probabilistic activation.

Consider a network of size *N*. Each node corresponds to an excitatory neuron, and spike propagation is approximated as a stochastic process at discrete time steps Δ*t*. If a neuron, described by the state variable *s*_*i*,*t*_ ∈ {0, 1}, is activated, it spikes (*s*_*i*,*t*_ = 1), and immediately returns to its resting state (*s*_*i*,*t*+1_ = 0) in the next time step, unless activated again. Furthermore, it may activate post-synaptic neurons *j* with probability *p*_*ij*,*t*_ = *w_ij_ α*_*j*,*t*_, where *w_ij_* ∈ {0, 1} indicates whether two neurons are synaptically connected, and *α*_*j*,*t*_ is a homeostatic scaling factor. In addition, each neuron receives network-independent external input at rate *h_i_*, representing external input from other brain areas, external stimuli, and importantly also spontaneous spiking of single neurons generated independently of pre-synaptic spikes (e.g. by spontaneous synaptic vesicle release [82, 83]). The uncorrelated external input homogeneously affects the network at rate *h_i_* = *h*, modeled as Poisson processes with an activation probability 1 − *e*^−*h*Δ*t*^ ≃ *h*Δ*t*.

This model can be treated in the framework of a branching process [77], a linear process with a characteristic autocorrelation time *τ* (see below). The population activity is characterized by the total number of spiking neurons,
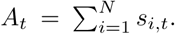
Each spike at time *t* generates on average *m* postsynaptic spikes at time *t* + 1 such that on average 𝔼(*A*_*t*+1_|*A_t_*) = *mA_t_*, where *m* is called *branching parameter*. The branching parameter can be defined for each neuron individually: neuron *i* activates on average

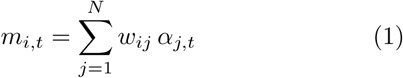

of its post-synaptic neurons [84]. This local branching parameter *m*_*i*,*t*_ thus quantifies the impact of a spike in neuron *i* on the network. The network average (denoted in the following with a bar) of *m*_*i*,*t*_ generates the (time-dependent) network branching parameter [65]

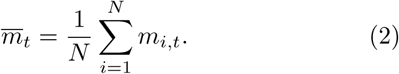

The external input generates on average *N h*Δ*t* additional spikes per time step, resulting in a driven branching process [85, 86]. The expected activity at time *t* + 1 is then 𝔼(*A*_*t*+1_|*A_t_*) = *mA_t_* + *N h*Δ*t*. For *m* < 1 the process is called *subcritical*, meaning that individual cascades of events will eventually die out over time. In this case, the temporal average (denoted in the following as 〈·〉) of network activity *A_t_* converges to a stationary distribution with average activity

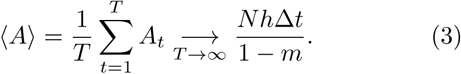

Considering a homogeneous neural spike rate *r_i_* = *r* = 〈*A*〉/*N* Δ*t* this implies

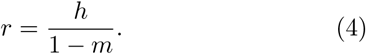

A constant mean spike rate *r*, which can be considered a biological constraint, is thus realized by adjusting either *m* ∈ [0, 1) or *h* ∈ [0, ∞).

The subcritical branching process (*m* < 1) is stationary with the autocorrelation function *C*(*l*) = *m^l^*. The autocorrelation time can be identified by comparing with an exponential decay *C*(*l*) = *e*^−*l*Δ*t*/*τ*^, yielding [80]

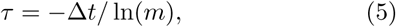

which diverges as *m* → 1. At this divergence (*m* = 1) the process is *critical* and the activity *A_t_* grows linearly in time with rate *h*. At criticality, assuming *h* → 0, the number of events *s* in an avalanche triggered by a single seed event, is distributed according to a power law *P* (*s*) ~ *s*^−3/2^ [77]. For a non-vanishing *h* in the sub-critical regime (*m* < 1), the avalanche-size distributions show a rapid decay, if they can be measured at all under persistent activity (Appendix B-2). Finally, for *m* > 1, the process is called *supercritical* and *A_t_* can in principle grow to infinity. For a finite network, this of course is not possible and will manifest in a peak of the avalanche-size distribution at large avalanche sizes.

For the computational model, we consider a network of *N* = 10^4^ neurons, which represents the size of *in vitro* cultures and *in vivo* cortical hypercolumns. The time step of the branching process has to reflect the causal signal propagation of the underlying physiological network.

Since realistic propagation times of action potentials from one neuron to the next range from 1 ms − 4 ms, we choose Δ*t* = 1 ms. We consider three network topologies:

#### a. Directed Erdős-Réenyi (ER) network

As a standard model of network topology, we consider a network with random directed connections. Each connection *w_ij_* = 1 is added with probability *p*_con_, excluding self-connections (*i*, *i*). Then, the degree distribution of outgoing as well as incoming connections follows a binomial distribution with average degree *k* = *p*_con_(*N* − 1) ≃ *p*_con_*N*. We require *p*_con_ > ln(*N*)/*N* to ensure that the graph is connected [87]. The connectivity matrix *w_ij_* is fixed throughout each simulation, such that averaging over simulations with different network realizations results in a quenched average. A cutout from an example graph is shown in Fig. 2**a**.

**FIG. 2.**
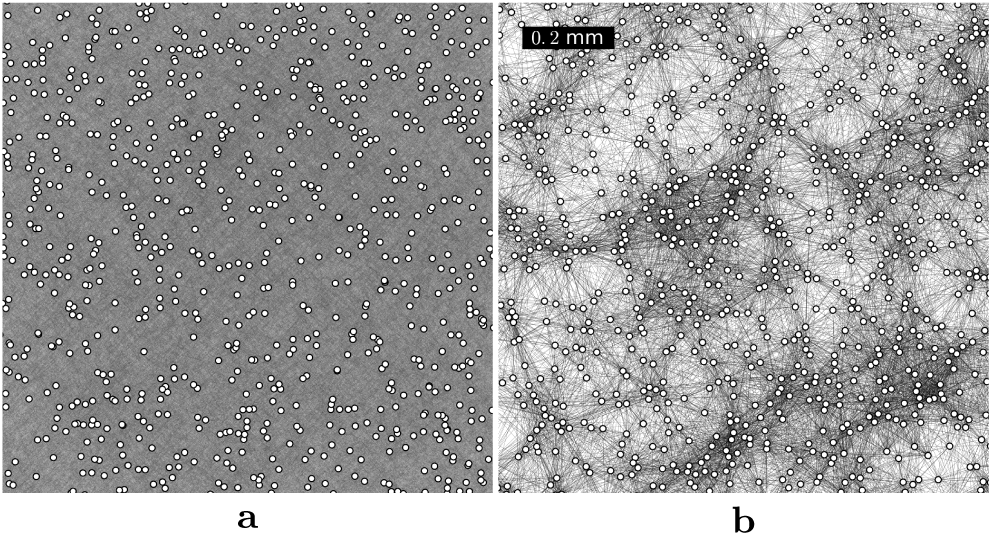
Cutouts of two random network topologies. (**a**) Subset of randomly spaced nodes in an Erdős-Rényi (ER) network with *p* = 10^−3^. Note that connections cross the window also from neurons outside of the field of view, such that single connections cannot be distinguished visually in the sketch. (**b**) 1.4 × 1.4mm^2^ subset of spatially-clustered (SC) topology generated by axonal-growth rules [5, 88].

#### b. Spatially-clustered (SC) network

In order to consider a more detailed topology with dominant short-range connections, we follow Orlandi *et al.* who developed a model based on experimental observations of *in vitro* cultures [5, 88]. Neural somata are randomly placed as hard discs with radius *R_s_* = 7.5 *μ*m, to account for the minimal distance between cell centers, on a 5 × 5 mm^2^ field. From each soma an axon grows into a random direction with a final length *l* given by a Rayleigh distribution
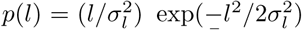
with *σ_l_* = 900 *μ*m and average axonal length *l̄* ≃ 1.1mm. The axonal growth is a semiflexible path with segments of size Δ*l* = 10 *μ*m and orientation drawn from a Gaussian distribution relative to the previous segment with *σ_θ_* = 15 °. A connection with another neuron is formed with probability 1/2 if the presynaptic axon intersects with the dendritic tree of a postsynaptic neuron [89]. The dendritic tree is modeled as a disc around the postsynaptic neuron with radius *R_d_* drawn from a Gaussian distribution with mean *R̄_d_* = 300 *μ*m and *σ_d_* = 20*μ*m. A cutout from an example graph is shown in Fig. 2**b**.

#### c. Annealed-average (AA) network

We consider in addition a network with *k* dynamically changing random connections (annealed average). The connections are distinguishable, exclude self-connections, and are redrawn every time step. This model approximates the otherwise numerically expensive fully connected network (ER with *p*_con_ = 1) with a global *m_t_* by choosing *α*_*j*,*t*_ = *m_t_*/*k*. In practice, we chose *k* = 4, which produces analogous dynamics to the fully-connected (*k* ≈ 10^4^) network as long as *m_t_* < 4.

Error bars are obtained as statistical errors from the fluctuations between independent simulations, which includes random network realizations {*w_ij_*} for ER and SC.

### B. Homeostatic plasticity

In our model, homeostatic plasticity is implemented as a negative feedback, which alters the synaptic strength on the level of the post-synaptic neuron (the homeostatic scaling factor *α*_*j*,*t*_) to reach a target neural firing rate
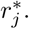
We consider a linear negative feedback with time constant *τ*_hp_, which depends solely on the (local) activity of the postsynaptic neuron *s*_*j*,*t*_

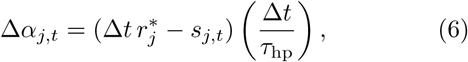

i.e., adapting a neuron’s synaptic strength does not rely on information about the population activity *A_t_*. Since *α*_*j*,*t*_ is a probability, we additionally demand *α*_*j*,*t*_ ≥ 0. Equation (6) considers homeostatic plasticity to directly couple to all postsynaptic synapses of any given neuron *j*. This can be implemented biologically as autonomous synaptic processes or somatic processes, such as translation and transcription. In order to further reduce complexity, we assume a uniform target rate
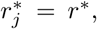
while in fact experiments show a broad (log-normal) spike-rate distribution [90, 91]. Preliminary tests for a log-normal target rate distribution in ER networks (*p*_con_ = 0.1) showed consistent results. In our simulations, we typically consider a biologically motivated target rate *r*^∗^ = 1 Hz and a homeostatic timescale of the order of an hour, *τ*_hp_ = 10^3^ s.

## IV. RESULTS

Including homeostatic plasticity in our model generates a broad range of dynamic states, depending on the external input. Figure 3 shows qualitatively representative results obtained for AA networks. For strong input (*h* = 𝓞(*r*^∗^)), the network organizes itself into a dynamic state where neural firing is solely driven by the input fluctuations, resembling an asynchronous-irregular state (green). Here, temporal and pairwise spike count cross-correlations approach zero, and the avalanche-size distribution matches the analytic solution for a Poisson process [92] shown as dashed lines. For weaker input (*h* < *r*^∗^) the system tunes itself towards fluctuating dynamics (orange-yellow). The average neural rate and sub-sampled avalanche-size distributions are qualitatively similar to reverberant *in vivo* dynamics with autocorrelation times of several hundred milliseconds (Fig. 1). In this regime, the temporal correlations increase when weakening the input, approaching close-to-critical dynamics, characterized by a power-law distribution *P* (*s*) = *s*^−3/2^ [77], at the lower end of the regime. Decreasing the input even further (*h* ≪ *r*^∗^) leads to bursting behavior, characterized by silent periods which are interrupted by strong bursts. These bursts are apparent as peak in the avalanche-size distribution at large avalanche sizes (purple-red). In this regime, the network steadily increases its synaptic strengths during silent periods until a spike initiates a large burst, which in turn decreases the synaptic strengths drastically, and so on (cf. Appendix C). This regime captures the qualitative features of bursting *in vitro* dynamics (Fig. 1).

**FIG. 3.**
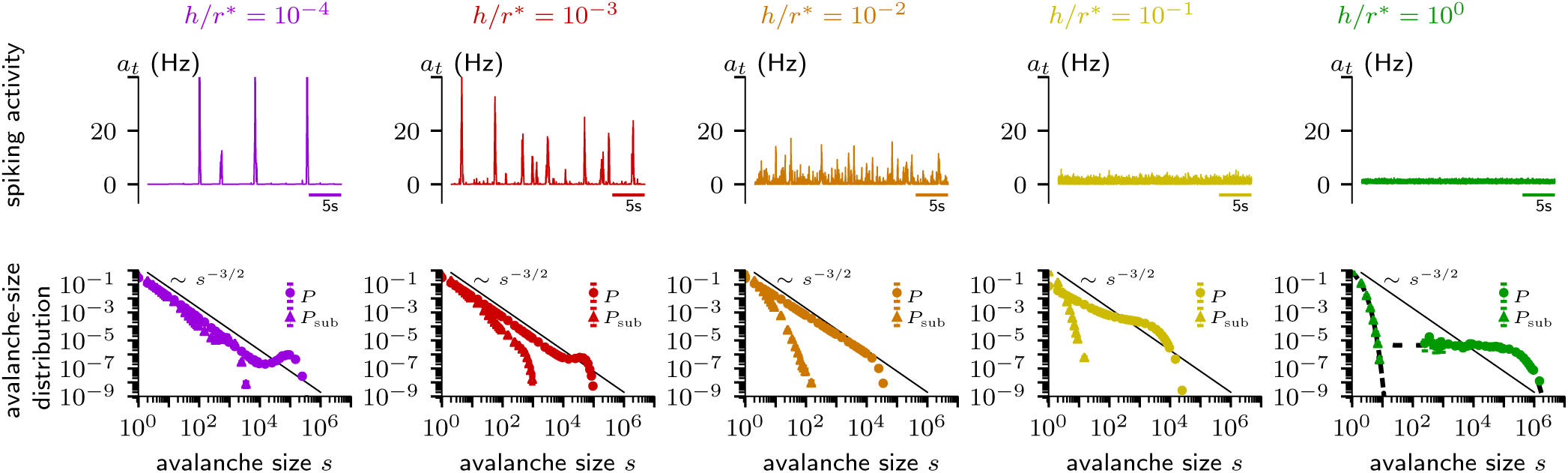
Homeostatic plasticity induces diverse dynamic states by regulating recurrent network interactions, mediating input rate *h* and target neural firing rate *r*^∗^. The annealed-average network reproduces bursting (*m* > 1, *h/r*^∗^ ≤ 10^−3^, purple-red), fluctuating (*m* ≈ 0.99,*h/r*^∗^ ≈ 10^−2^ and *m* ≈ 0.9, *h/r*^∗^ ≈ 10^−1^, orange-yellow), and irregular (*m* ≈ 0, *h/r*^∗^ = 1, green) dynamics. The top row shows examplary spiking activity *a_t_* = *A_t_/N* Δ*t* (Appendix B-1); the bottom row shows avalanche-size distributions *P* (*s*) (*n* = *N*, circles) and subsampled avalanche-size distributions *P*_sub_(*s*) (*n* = 100, triangles) averaged over 12 independent simulations (Appendix B-2). Solid lines show the power-law distribution *P* (*s*) ∝ *s*^−3/2^ [77], dashed lines show the analytical avalanche-size distribution of a Poisson process [92]. Parameters: *N* = 10^4^, *τ*_hp_ = 10^3^ s, *r*^∗^ = 1 Hz, Δ*t* = 1 ms.

In the following, we derive a quantitative description of the three regimes sketched above. To quantify the dynamic state, we consider the temporal average of the branching parameter *m* = 〈*m*〉, as well as the associated autocorrelation time *τ* of the population activity.

### A. Mean-field solution

If we assume that *τ*_hp_ is sufficiently large (i.e. slow homeostatic plasticity), then Δ*α_j_* ≈ 0 and the dynamics of the network is fully determined by the approximately constant branching parameter *m_t_* ≈ *m*. In this regime, (4) holds and combined with (5) and (6) we obtain the mean-field solution

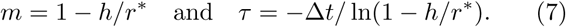

Hence, with decreasing input rate *h*, recurrent network activation (*m*) increases, i.e., perturbations cause a stronger network response and the autocorrelation time increases (Fig. 4, solid lines).

**FIG. 4.**
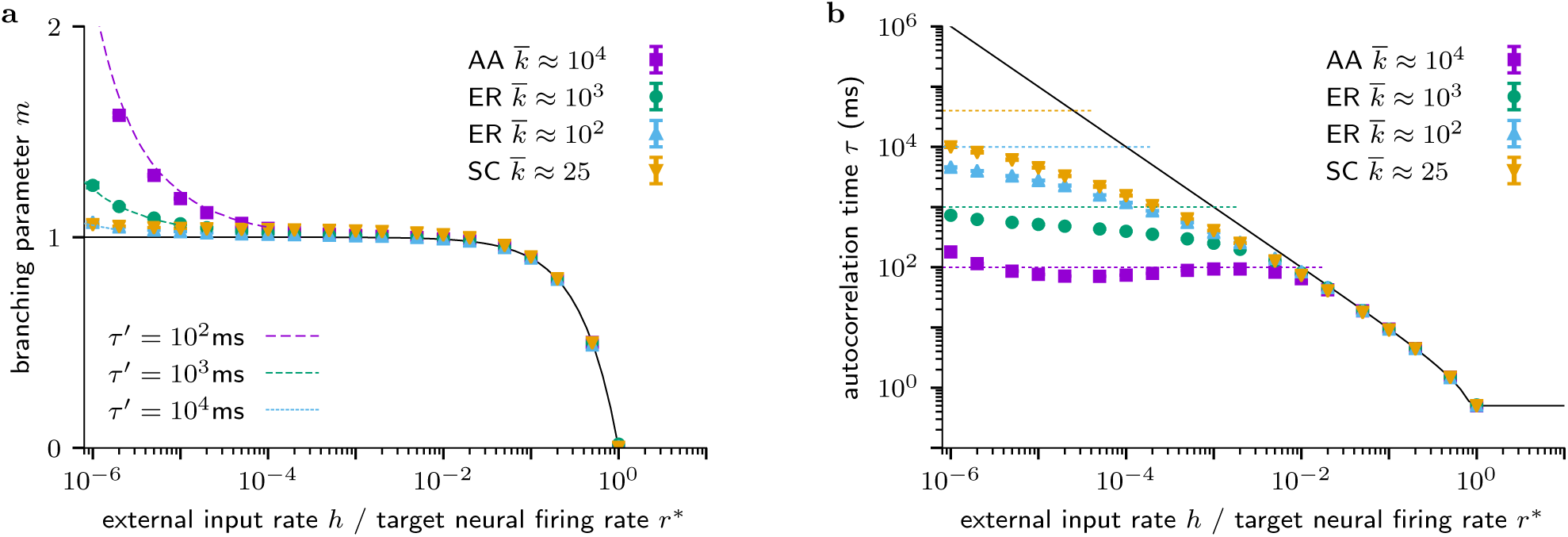
Quantitative distinction between dynamic states induced in neural networks of different topologies by homeostatic plasticity as a function of normalized input rate *h/r*^∗^. Data points are averages over 12 independent simulations (*N* = 10^4^, *τ_α_* = 10^3^ s, *r*^∗^ = 1 Hz, Δ*t* = 10^−3^ s) with connections generated according to annealed-average (AA), Erdős-Réenyi (ER) or spatially-clustered (SC) topologies with average number of connections *k*. Solid lines show the mean-field solution (7), dashed lines represent (semi-analytical) approximations of the bursting regime. (**a**) Branching parameter *m* = 〈*m*〉 varies from irregular (*m* ≈ 0), to fluctuating (*m* ⪅ 1), to bursting (*m* > 1) dynamics. The behavior in the bursting regime strongly depends on the network timescale *τ′* = *τ*_hp_/*k*. (**b**) Integrated autocorrelation time of the network population activity (Appendix B-3) shows a crossover from irregular [*τ* = 𝓞(Δ*t*)], over fluctuating [*τ* = −Δ*t*/ ln(1 − *h/r*^∗^)] to bursting (*τ* ≈ *τ′*) dynamics.

In the light of this mean-field solution, we discriminate the three characteristic regimes as follows. First, we define the *input*-*driven regime* by *m* ≤ 0.5 and *τ* ≈ Δ*t*. Here, the network activity is dominated by input (*h* = 𝓞(*r*^∗^)), and thus the dynamics follows the input statistics and becomes irregular. Second, we define the *fluctuating regime* for 0.5 < *m* < 1 with a non-vanishing but finite autocorrelation time Δ*t* < *τ* < ∞. Here, the network maintains and amplifies input as recurrently generated fluctuations. In these two regimes the mean-field solution (7) matches numerical data on different network topologies (Fig. 4). Third, the mean-field solution predicts that in the limit *h* → 0 the dynamics become critical with divergent autocorrelation time (*m* → 1, *τ* → ∞). However, we observe a clear deviation from the mean-field solution, which defines the *bursting regime* with *m* > 1 and a finite autocorrelation time, as discussed below.

### B. Bursting regime

Deviations from the mean-field solution (7) emerge when the assumption of “sufficiently large *τ*_hp_” breaks down. We will derive a bound for *τ*_hp_, below which the (rapid) homeostatic feedback causes notable changes of the network branching parameter *m_t_* around its mean *m* = 〈*m*〉, which in turn jeopardize the stability of the network dynamics.

To estimate the change of the network branching parameter, we first consider the change in local branching parameter Δ_*mi*,*t*_, which depends on each neurons out-degree
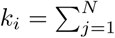
*w_ij_* and is given by

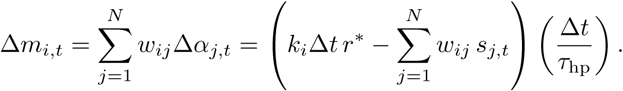

On the network level, we make the assumption that the state of each neuron is approximated by the network average *s*_*i*,*t*_ ≈ *A_t_/N*, such that 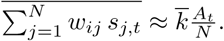
Then, the change in network branching parameter can be approximated as

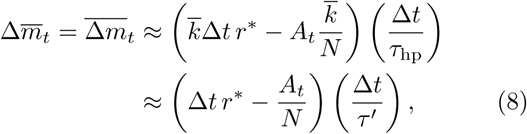

where we have introduced an effective homeostatic network timescale *τ′* = *τ*_hp_*/k*, for which (8) recovers the form of (6). Using *τ′* allows one to semi-analytically approximate the deviation of *m* from the mean-field solution (Fig. 4**a**, dashed lines, and Appendix C).

We next show that the stability of network dynamics requires the autocorrelation time of the dynamic process *τ* to be smaller than the timescale of homeostasis *τ′*. Stability demands that the homeostatic change in autocorrelation time Δ*τ* is small compared to the autocorrelation time itself, i.e., Δ*τ* ≪ *τ*. We approximate Δ*τ* by error propagation in (5), yielding

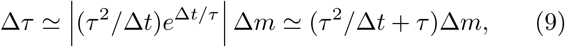

where we expanded the exponential for small Δ*t/τ*. For large *τ*, the leading term in (9) dominates and inserting (8) yields Δ*τ* ≃ |Δ*t r*^∗^ − *A_t_/N*| (*τ*^2^/*τ′*). Thus, the dynamics can be described as a stationary branching process (mean-field solution) only as long as

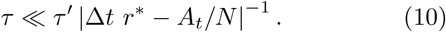

Violation of (10) results in bursting behavior (Figs. 3 & 7). For *A_t_* = 𝓞(*N*) the right hand side of (10) is minimal, because Δ*t r*^∗^ ≪ 1, which implies a maximal attainable autocorrelation time *τ* ≃ *τ′* = *τ*_hp_*/k*. This is in perfect agreement with the saturation of measured autocorrelation time in the bursting regime (Fig. 4**b**, dashed lines).

The transition from the fluctuating to the bursting regime occurs when the mean-field solution (7) equals the maximal attainable autocorrelation time, i.e., *τ* = −Δ*t*/ ln(1 − *h/r*^∗^) ≈ *τ′*. Hence, the transition occurs at *h/r*^∗^ ≈ 1 − *e*^−Δ*t*/*τ′*^ ≈ Δ*t*/*τ′*. For even lower input rate, the dynamics become more and more bursty, and the avalanche-size distribution exhibits a peak at large avalanche sizes (Fig. 3 for *h/r*^∗^ < 10^−2^, where *τ′* = 10^2^ ms, Δ*t* = 1 ms). At the transition, the dynamics can be considered close-to-critical, because the (fully sampled) avalanche-size distribution is closest to a power-law with exponent ^−3/2^.

### C. Distributions of spiking activity

The different dynamical regimes imply characteristic distributions of neural network activity *P*(*a_t_*). Figure 5 shows an example of *P*(*a_t_*) for ER networks with *p*_con_ = 10^−2^, where the transition from fluctuating to bursting dynamics is expected for *h/r*^∗^ ≈ Δ*t*/*τ′* = 10^−4^. In the irregular regime (green) *P*(*a_t_*) is a unimodal distribution. In the fluctuating regime (yellow-red), the peak in *P*(*a_t_*) shifts towards quiescence and the distribution develops a power-law tail with exponential cutoff, expected for a critical branching process. In the bursting regime (purple-blue), *P*(*a_t_*) is a bimodal distribution, reflecting network changes between quiescence and bursty activity. The position and sharpness of the high-activity maximum depend on the network connectivity and hence the heterogeneity in the single-neuron input.

**FIG. 5.**
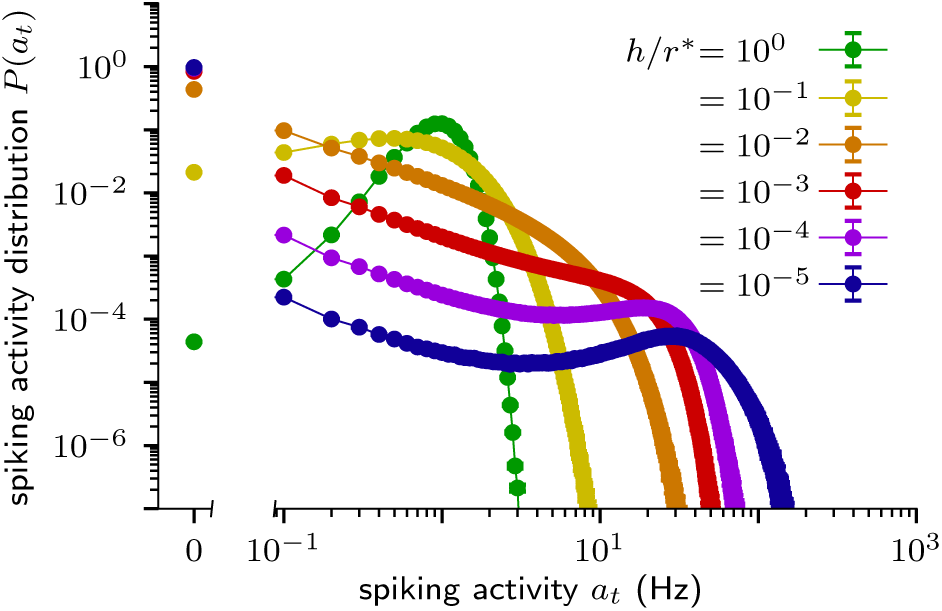
Distribution of spiking activity in weakly connected Erdős-Réenyi networks (*p*_con_ = 10^−2^, Δ*t* = 1 ms, *r*^∗^ = 1 Hz, *τ′* = 10^4^ ms) averaged over 12 independent simulations. For irregular dynamics (*h/r*^∗^ ≈ 10^0^) the distribution is clearly unimodal. For fluctuating dynamics (10^−4^ < *h/r*^∗^ < 10^0^) the distribution broadens and shifts the maximum towards quiescence. In addition towards the lower bound of the regime, the distribution develops a power-law tail with an exponential cutoff. At the crossover to bursting dynamics (*h/r*^∗^ ≈ 10^−4^) the distribution becomes bimodal.

### D. Reproducing experimental results

Using the insight from our theory, we can reproduce experimental results. Spiking activity recorded *in vivo* resembles dynamics of the fluctuating regime. In this regime, the dynamic state is consistent for all topologies we considered (Fig. 4). Therefore, already a branching network on an AA topology suffices to quantitatively reproduce the avalanche-size distributions by matching model parameters with experimentally accessible estimates (Fig. 1 dashed lines). To match the branching network to recordings from cat V1 and rat CA1, we first estimated the spike rate *r* and autocorrelation time *τ* from the recordings of spiking activity [80]; we then chose biologically plausible parameters for the network size *N*, the homeostatic timescale *τ*_hp_, as well as the simulation time step Δ*t*; and finally derived the external input *h* using (7) (for details see Appendix B-4). The resulting subsampled avalanche-size distributions are in astonishing agreement with the experimental results, given the simplicity of our approach. Close inspection of the avalanche-size distribution for rat CA1 recordings still reveals small deviations from our model results. The deviations can be attributed to theta-oscillations in hippocampus, which result in subleading oscillations on an exponentially decaying autocorrelation function [80]. While this justifies our approach to consider a single dominant autocorrelation time, theta oscillations slightly decorrelate the activity at short times and thereby foster premature termination of avalanches. Thus, the tail in the avalanche-size distribution is slightly shifted to smaller avalanche sizes (Fig. 1).

The *in vitro* results are qualitatively well matched by simulations in the bursting regime, with avalanche-size distributions showing a characteristic peak at large avalanche sizes (Fig. 3). It is difficult to quantitatively match a model to the data, because a number of parameters can only be assessed imprecisely. Most importantly, the autocorrelation time in the burst regime is not informative about the external input rate *h* and depends on the average number of connections (Fig. 4). Likewise, the time-dependence of the branching parameter *m_t_* cannot be assessed directly. Finally, system size and topology impact the network dynamics more strongly in this regime than in the fluctuating or input-driven regime. This yields a family of avalanche-size distributions with similar qualitative characteristics but differences in precise location and shape of the peak at large sizes.

## V. DISCUSSION

We propose the interplay of external input rate and target spike rate, mediated by homeostatic plasticity, as a neural mechanism for self-organization into different dynamic states (cf. sketch in Fig. 6). Using the framework of a branching process, we disentangled the recurrent network dynamics from the external input (e.g. input from other brain areas, external stimuli and spontaneous spiking of individual neurons). Our mean-field solutions, complemented by numeric results for generic spiking neural networks, show that for high input the network organizes into an input-driven state, while for decreasing input the recurrent interactions are strengthened, leading to a regime of fluctuating dynamics, resembling the reverberating dynamics observed *in vivo*. Decreasing the input further induces bursting behavior, known from *in vitro* recordings, due to a competition of timescales between homeostatic plasticity and the autocorrelation of population activity. Thereby our framework proposes a generic mechanism to explain the prominent differences between *in vivo* and *in vitro* dynamics.

**FIG. 6.**
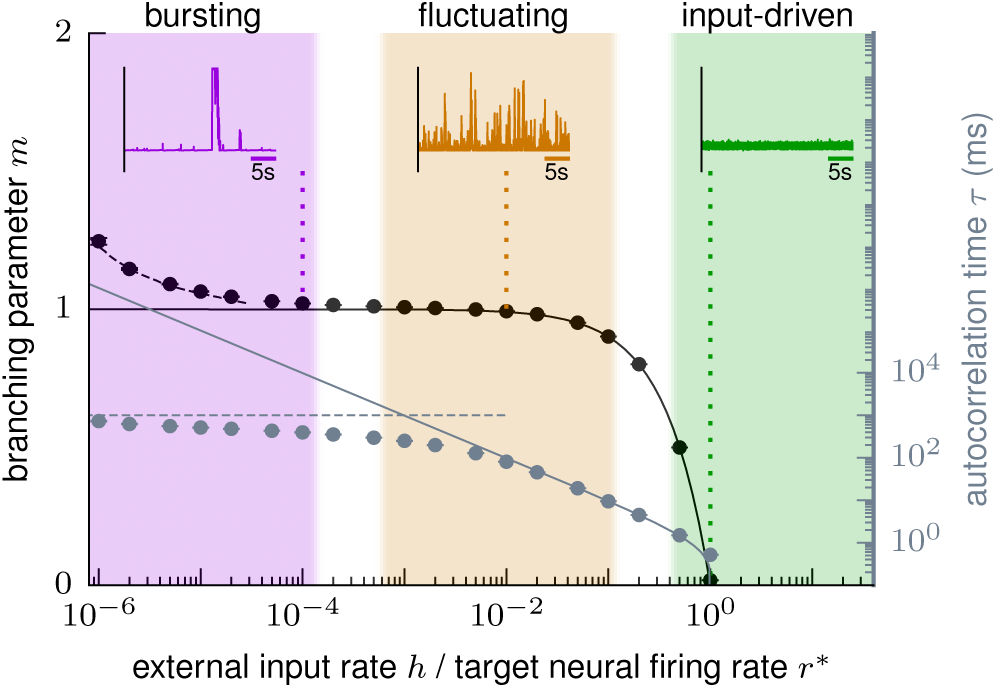
Sketch of bursting, fluctuating and input-driven net-
work states, classified by the branching parameter and the autocorrelation time. We propose (solid lines) that homeostatic plasticity tunes the dynamic state depending on the ratio of external input rate (including spontaneous neural firing) and target neural firing rate. Data points and example activity traces stem from Erdős-Rényi networks (*N* = 10^4^, *p* = 10^−1^, *τ*_hp_ = 10^3^ s). In the bursting regime, the homeostatic timescale *τ*_hp_ influences the resulting dynamics (dashed lines).

Our theory suggests that also differences within the collective dynamic state observed *in vivo* can be explained by considering differences in input strength. For cortex, it was shown that layer 2/3 exhibits critical dynamics [17] and presumably deeper layers show reverberating dynamics [80, 81]. We propose that this can be caused by different input strength: layer 2/3 is more recurrently connected, while layer 4 is the main target of thalamic input [93], hence receiving the stronger input. The dynamic state varies also across different cortical areas, where autocorrelation times of network activity reflect a hierarchical organization [8, 94]: Cortical areas associated with higher-order function show a larger autocorrelation time. In the light of our results, a larger autocorrelation time implies less afferent input for the area in question. The hierarchical organization is further supported by our analysis of spiking activity *in vivo* (Fig. 1): the autocorrelation times in visual cortex (*τ* ≈ 0.2 s) and hippocampus (*τ* ≈ 2 s) precisely reflect that visual cortex is almost at the bottom, whereas hippocampus is at the top of the hierarchy of visual processing [95].

Our theory provides an approach to experimentally infer the fraction of spikes generated recurrently within a network and generated by external input. For an average spike rate *r*, equation (7) implies *h/r* = (1 – *e*^−Δ*t*/τ^). The external input rate can then be directly calculated from the autocorrelation time and by assuming a biologically plausible signal-propagation time, e.g., Δ*t* ≈ 4 ms. We estimate for recordings from visual cortex in mildly anesthetized cat that about 2% of the network activity is generated by the input, whereas the majority of 98% are generated recurrently within the network. From autocorrelation times measured across the cortical hierarchy (50 ms to 350 ms) in macaque monkey [8], the fraction of spikes generated by external input decreases from ~ 8 % to ~ 1 % from lower to higher cortical areas. This is consistent with perturbation experiments in rat barrel cortex, where after triggering an extra spike the decay time of population rate was at least 50 ms [96] indicating at most about 8% external input (for a detailed discussion see also Ref. [97]). Last, experiments on visual cortex of awake mice directly after thalamic silencing found a decay time of *τ* = 12(1) ms [98], from which we would estimate about 70% recurrent activation. This is in perfect agreement with the experimentally measured 72(6)% of recurrent activation in the same study. This result thus validates our derived relation between *h/r* and *τ*.

One can interpret our findings in the light of up and down states [18, 19, 65, 99]. Because the membrane potential was found to correlate with network activity [6, 20], our results for the distribution of spiking activity in the bursting regime may correspond to the bimodal distributions of membrane potentials during up and down states (Fig 5). It has already been shown that negative feedback can stabilize up and down states [65, 99]. In our theory, negative feedback leads to similar results in the low-input regime. Moreover, we predict that decreasing network input further, prolongs the quiescent periods or down states.

Our theory unifies previous numerical approaches of self-organization in neural networks, which typically considered a negative feedback mechanism but made very different choices on a (fixed) network input. For example, bursting dynamics have been generated by homeostatic build-up upon loss of network input [43] or by self-organized supercriticality through dynamic neuronal gain [100]. Adding weak input, self-organized criticality [101, 102] has been achieved by local rewiring [55–57] and synaptic depression [51, 52, 58–63]. In contrast, asynchronous-irregular network activity typically requires comparably strong input, assuming a balanced state [47, 103, 104], and a self-organized AI network state can be promoted by inhibitory plasticity [53, 54]. While all these studies provide mechanisms of self-organization to one particular dynamic state, our theory highlights the role of input in combination with a negative feedback [40, 47–49] and provides a unifying mechanism of self-organization covering bursting, fluctuating and irregular dynamics.

From a broader perspective, we characterized driven systems with a negative feedback as a function of the input rate. The negative feedback compensates the input by regulating the system’s self-activation to achieve a target activity. In this light of control theory, the bursting regime can be understood as resonances in a feedback loop, where feedback dynamics are faster than system dynamics (cf. [105]). This qualitative picture should remain valid for other connected graphs subject to external input with spatial and temporal correlations. In this case, however, we expect more complex network responses than predicted by our mean-field theory, which assumes self-averaging random networks subject to uncorrelated input.

Our results suggest that homeostatic plasticity may be exploited in experiments to generate *in vivo*-like dynamics in a controlled *in vitro* setup, in particular to abolish the ubiquitous bursts *in vitro*. Previous attempts to reduce bursting *in vitro* [106] and in model-systems of epilepsy [107–110] used short-term electrical and optical stimulation to attain temporal reduction in bursting. Alternatively, one can reduce bursting pharmacologically or by changing the calcium level, however, typically at the cost of changing single-neuron properties [111–113]. We propose a different approach, namely applying weak, global, long-term stimulation. Mediated by homeostasis, the stimulation should alter the effective synaptic strength, and thereby the dynamic state while preserving single-neuron dynamics [114]. In particular, we predict that inducing in every neuron additional spikes with *h* = 𝓞(0.01 Hz) is sufficient to abolish the ubiquitous bursts *in vitro* and render the dynamics *in vivo*-like instead. If verified, this approach promises completely novel paths for drug studies. By establishing *in vivo*-like dynamics *in vitro*, fine differences between neurological disorders, which are otherwise masked by the ubiquitous bursts, can be readily identified. Altogether this would present a comparably cost-efficient, high-throughput, and well-accessible drug assay with largely increased sensitivity.

## ACKNOWLEDGMENTS

We would like to thank Manuel Schottdorf and Andreas Neef, as well as Raoul-Martin Memmesheimer and Sven Goedeke, for stimulating discussions. We are grateful for careful proofreading from Roman Engelhardt, Joãao Pinheiro Neto, and Conor Heins. All authors acknowledge funding by the Max Planck Society. JZ and VP received financial support from the German Ministry of Education and Research (BMBF) via the Bernstein Center for Computational Neuroscience (BCCN) Göttingen under Grant No. 01GQ1005B. JW was financially supported by Gertrud-Reemtsma-Stiftung and Physics-to-Medicine Initiative Göttingen (ZN3108) LM des Niedersächsischen Vorab.

## Appendix A: Experimental details

### 1. Dissociated dense cultures of cortical rat neurons

The spike-time data from dissociated cortical rat neurons of mature dense cultures was recorded by Wagenaar *et al.* [4] and was obtained freely online [72]. The experimental setup uses multi-electrode-arrays (MEA) with *n* = 59 electrodes. Cortical cells were obtained from dissecting the anterior part of the cortex of Wistar rat embryos (E18), including somatosensory, motor, and association areas. For details, we refer to [4]. Measurements were performed every day *in vitro* (DIV). We here focus on the dense case with 50 000 cells plated initially with a density of 2.5(1.5) × 10^3^ cells/mm^2^ at 1 DIV, which is compatible with standard *in vitro* experiments in the field that claim to observe critical dynamic behavior. We selected the representative recordings 8-2-34 (exp 1) and 7-2-35 (exp 2) at mature age (34/35 DIVs) for Fig. 1.

### 2. Rat hippocampus

The spiking data from rats were recorded by Mizuseki *et al.* [73, 74] with experimental protocols approved by the Institutional Animal Care and Use Committee of Rutgers University. The data was obtained from the NSF-founded CRCNS data sharing website [74]. The spikes were recorded in CA1 of the right dorsal hippocampus during an open field task. Specifically, we used the data set ec013.527 with sorted spikes from 4 shanks with *n* = 31 channels. For details we refer to Refs. [73, 74].

### 3. Primary visual cat cortex

The spiking data from cats were recorded by Tim Blanche in the laboratory of Nicholas Swindale, University of British Columbia, in accordance with guidelines established by the Canadian Council for Animal Care [75, 76]. The data was obtained from the NSF-founded CRCNS data sharing website [76]. Specifically, we used the data set pvc3 with recordings of *n* = 50 sorted single units [75] in area 18. For details we refer to Refs. [75, 76]. We confined ourselves to the experiments where no stimuli were presented such that spikes reflect the spontaneous activity in the visual cortex of mildly anesthetized cats. In order to circumvent potential non-stationarities at the beginning and end of the recording, we omitted the initial 25 s and stopped after 320 s of recording [80].

## Appendix B: Analysis details

### 1. Spiking activity

In order to present the spiking activity over time, we partition the time axis of experimental or numerical data into discrete bins of size Δ*t*. For the time-discrete simulations the time bin naturally matches the time step. For experimental data we set Δ*t* = 4 ms. In each time bin we count the total number of spikes *A_t_* and normalize with the number of neurons *N* to obtain the average spiking activity *a_t_* = *A_t_/N*Δ*t*. Note that experimental preparations were inevitably subsampled, as spikes were recorded only from a small number of all neurons.

### 2. Avalanche-size distribution

We define the avalanche size *s* as the number of spikes enclosed along the discrete time axis by bins with zero activity [7]. To test for criticality in terms of a branching process, one compares *P* (*s*) to the expected *P* (*s*) ~ *s*^−3/2^. This is a valid approach in the limit *h* → 0, where avalanches can be clearly identified, and for fully sampled systems [81]. However, experiments are limited to record only from *n* out of *N* neurons. As a result, the distributions for subsampled activity *P*_sub_(*s*) differ due to subsampling bias [15, 16]. Therefore, we numerically measure both full (*n* = *N*) and subsampled (*n* < *N*) avalanche-size distributions to qualitatively compare *P*(*s*) to the theory and *P*_sub_(*s*) to experimental data.

### 3. Integrated autocorrelation time

We measure the autocorrelation time of spiking activity *a_t_* in terms of the integrated autocorrelation time *τ*_int_, for details see, e.g., Ref. [115]. In brief, we sum over the normalized autocorrelation function *C*(*l*) = Cov[*a_t_*, *a*_*t*+*l*_]/Var[*a_t_*] until the sum converges. Following conventions, we define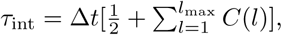 where *l*_max_ is self-consistently obtained as the minimal *l*_max_ > 6 *τ*_int_(*l*_max_).

### 4. Reproducing experimental results

We use a branching network with AA topology subject to homeostatic plasticity to quantitatively reproduce *in vivo* sub-sampled avalanche-size distributions. We chose networks of size *N* = 10^4^ with sufficiently large homeostatic timescale *τ*_hp_ = 10^5^ s. The following model parameters can be obtained from experimentally measured values: In the chosen recordings, we measured the average rate (*r*_cat_ ≈ 7 Hz and *r*_rat_ ≈ 11 Hz) as well as the subsampling corrected branching parameter [80] (*m*_cat_ ≈ 0.98 and *m*_rat_ ≈ 0.997 for Δ*t* = 4 ms). In fact, the branching parameter is not suitable to identify the input rate via (7), because it refers to a process in discrete time steps. Since we are treating a continuous process, the invariant quantity is the autocorrelation time (*τ*_cat_ ≈ 0.2 s and *τ*_rat_ ≈ 1.6 s). According to our theory, we can then calculate the input rate per neuron *h* = (1–exp(–Δ*t/τ*))*r*. In order to avoid convergence effects, we need to choose a sufficiently small time step Δ*t* = 1 ms of signal propagation (resulting in *h*_cat_ ≈ 3.5 × 10^−2^ Hz and *h*_rat_ ≈ 5.5 × 10^−3^ Hz), while we record in time bins of 4 ms to match the analysis of the experiments. Subsampled avalanche-size distributions are estimated by randomly choosing *n* < *N* neurons, where we approximated *n* by the number of electrodes or channels (*n*_cat_ = 50 and *n*_rat_ = 31).

## Appendix C: Approximating the dynamic state in the bursting regime

We showed in Sec. IV that decreasing the external input to recurrent networks with homeostatic plasticity leads to bursting behavior (Fig. 7a). This is directly related to the network branching parameter *m_t_* = *m_t_* no longer showing small fluctuations around the predicted value but instead exhibiting a prominent saw-tooth pattern (Fig. 7b), a hallmark of the homeostatic buildup in the long pauses with no input.

**FIG. 7.**
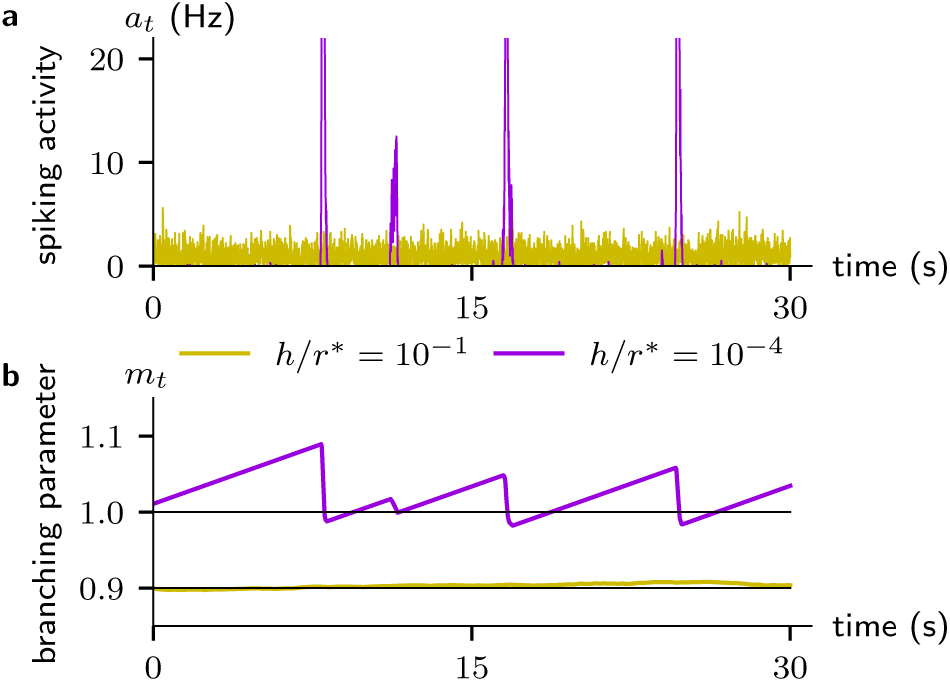
Temporal fluctuations in an annealed-average network with homeostatic plasticity subject to different external input rates. (**a**) Spiking activity *a_t_* shows small fluctuations for large input rates (yellow) and bursts for small input rates (purple), cf. Fig. 3. (**b**) Branching parameter fluctuates around predicted value (black horizontal lines) and develops distinct saw-tooth pattern for small input rates.

We here show a semi-analytical approximation of the network branching parameter in the bursting regime. For sufficiently small external input we may assume separation of timescales, i.e., every externally induced spike drives one avalanche with periods of silence in between. Let us first consider the periods of silence, i.e., no activity per site. This holds during the entire growth period

*T* such that (8) yields

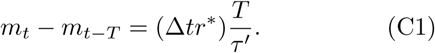

The situation becomes more involved within the bursts, where the behavior of *m_t_* is nonlinear. Consider an external spike that triggers an avalanche at *t* = *s* which ends at *t* = *e*. Due to the separation of timescales we can assume *A_s_* = 1. There are two possible scenarios: (i) The avalanche dies out before a burst can develop and (ii) the input triggers a proper burst with a macroscopic activation.

We first estimate the probability that an avalanche dies out before a burst develops. For *τ*_hp_ ≫ Δ*t* we approximate *m_t_* ≈ *m_s_* = const. Then, the probability of ultimate extinction *θ* can be calculated as the solution of *θ* = Π(*θ*) with Π(*θ*) the probability generating function [77]. In the onset phase, the branching process is described by a Poisson process per event with mean *m_s_*, such that Π(*θ*) = *e*^−*m_s_* (1–*θ*)^. We are thus looking for a solution of

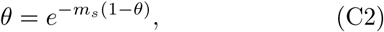

which can be rewritten to

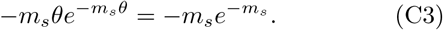

We identify the Lambert-W function *W*(*z*)*e*^*W*(*z*)^ = *z* [116] with *W*(*z*) = –*m_s_θ* = *W*(–*m_s_e*^−*m_s_*^) and find for the probability that no burst develops

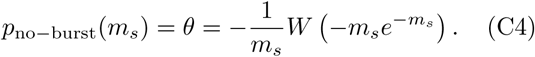

If a proper burst develops, the strong activity diminishes *m_t_* until the burst dies out again. We cannot analytically estimate the branching parameter *m_e_* after burst end, but we can use a deterministic numerical approximation to obtain *m_e_*(*m_s_*). Instead of stochastically generating new (discrete) events according to some distribution *P*(*m_t_*) with average *m_t_*, we approximate the branching process as deterministic (continuous) evolution *A*_*t*+1_ = *m_t_A_t_*. For a finite network, we need to consider convergence effects when one neuron is activated by two or more neurons at the same time. In the absence of external input, this introduces for an AA (i.e. approximating fully connected) network the activity-dependent branching parameter [117]

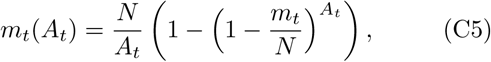

which we need to consider for the activity propagation within the burst, i.e., *A*_*t*+1_ = *m_t_*(*A_t_*)*A_t_*. In addition, we introduce an upper bound *A_t_* ≤ *N*. The upper limit on *A_t_* puts a lower bound on Δ*m_t_* according to (8) and thus extends the duration of avalanches. Evolving *m*_*t*+1_ = *m_t_* + (Δ*tr*^∗^ – *A_t_*/*N*)(Δ*t*/*τ′*), with *m*_*t*+1_ ≥ 0, we iterate until *A_e_* < 1. This is a quick and numerically robust iterative scheme to estimate *m_e_*(*m_s_*).

Putting everything together, we numerically approximate the average network branching parameter *m* under homeostatic plasticity in the bursting regime of low external input for an AA network. For this, we sample the external spikes (drive) as 10^4^ inter-drive intervals *T_s_* from an exponential distribution *P*(*T*) = (1/*hN*)*e*^−*T*/*hN*^, corresponding to *N* Poisson processes with rate *h*. The remaining part can be interpreted as an event-based sampling with approximate transformations: Starting with *m*_0_ = 0, we evolve *m_t_* for each inter-drive interval *T_s_* according to (C1). If *m_t_* > 1, we keep *m_t_* with probability *p*_no–burst_(*m_t_*) or else initiate a burst by setting *m_t_* = *m_e_* (*m_t_*). Afterwards we continue evolving *m_t_*.

## Appendix D: Characteristic duration of inter-burst-intervals in burst regime

In the bursting regime of low external input, the spiking activity suggests a characteristic time between bursts. In order to test for periodicity, we analyzed the distribution of inter-burst-intervals (IBI), where intervals are measured as the time between two consecutive burst onsets, defined as a spiking activity *a_t_* > 20*r*^∗^. We find (Fig. 8) that large IBI are suppressed by the exponentially distributed inter-drive intervals (dashed lines), while short IBI are suppressed by the probability *p*_no–burst_(*m*) that a given external spike does not trigger a proper burst (Appendix C). This gives rise to a characteristic duration of inter-burst-intervals in the burst regime, although the dynamics are not strictly periodic.

**FIG. 8.**
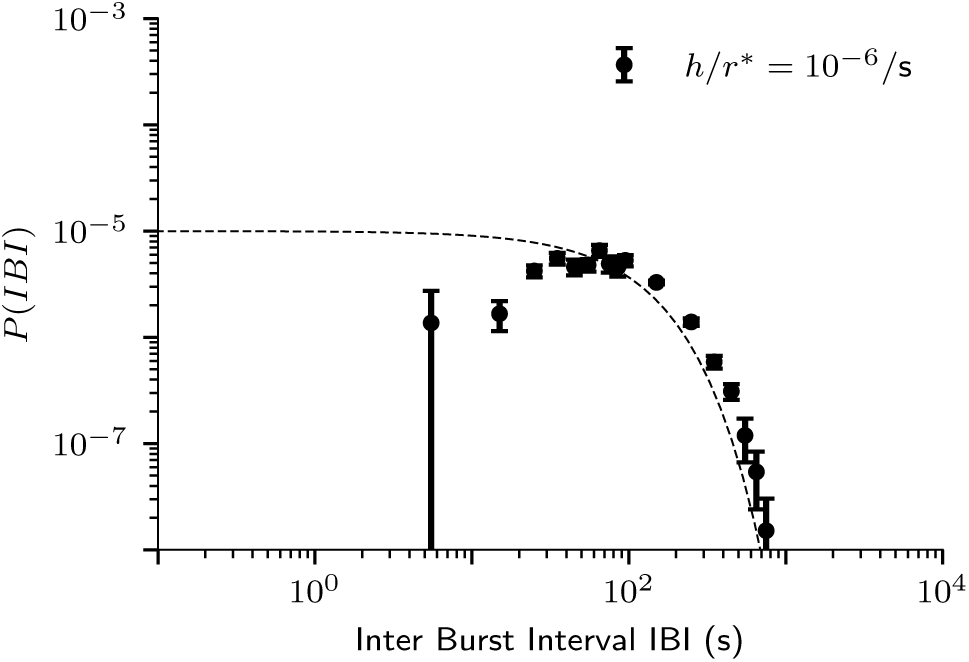
Inter-burst-interval (IBI) distribution for annealed-average networks averaged over 12 independent simulations. Intervals are measured as times between proper burst onsets (*a_t_* > 20*r*^∗^). Dashed lines show the exponential inter-spike-interval distribution of the Poisson external drive.

